# Simply cut out – Combining CRISPR/Cas9 RNPs and transiently selected telomere vectors for marker free-gene deletion in *Trichoderma atroviride*

**DOI:** 10.1101/2025.05.06.652419

**Authors:** Mario Gründlinger, Chiara Ellensohn, Leo Drechsel, Ulrike Schreiner, Siebe Pierson, Clara Baldin, Susanne Zeilinger

## Abstract

*Trichoderma atroviride* is a well-known mycoparasitic fungus widely used for the biological control of fungal plant pathogens. Expanding its genetic toolbox is essential to facilitate efficient genetic manipulation, including successive transformations and multiple or reusable selection markers for consecutive gene deletions. We applied CRISPR/Cas9 via ribonucleoprotein (RNP) complexes for gene editing in *T. atroviride* and successfully deleted three target genes, i.e., *pks4* (involved in spore pigment production), *pyr4* (pyrimidine biosynthesis), and *pex5* (receptor for peroxisomal matrix protein import). Although double-strand breaks induced by Cas9 can be repaired via homology-directed repair (HDR), using donor templates, the most effective gene deletions in our case were achieved via non-homologous end joining (NHEJ), by co-transforming a transiently stable telomere vector carrying the hygromycin-resistance gene (*hph*), which was rapidly lost under non-selective conditions. This strategy promoted NHEJ repair and resulted in the efficient deletion of open reading frames between two Cas9 target sites. Our results demonstrate that combining CRISPR/Cas9 RNP delivery with transient telomere vectors provides a fast and reliable method for marker-free gene deletion and vector recycling in *T. atroviride*, advancing the reverse genetic toolkit available for this important biocontrol fungus.

## 2 Introduction

*Trichoderma* fungi are widespread filamentous ascomycetes and several members are well known as enzyme producers in the biotechnological industry or as biological control agents and biofungicides in plant disease control [1]. The latter is based on the mycoparasitic lifestyle of *Trichoderma* that enables these fungi to parasitize a broad variety of economically relevant phytopathogens while at the same time shielding plant roots against pathogen attack and stimulating plant growth and defence [2, 3]. *Trichoderma atroviride* is a strong and commercially applied biocontrol agent and has been established as a model mycoparasite [4]. Despite having been studied and exploited for decades, many of the over 11,000 genes and gene products encoded in the *T. atroviride* genome still are functionally uncharacterized.

Reverse genetics is commonly applied for the characterization of gene function in fungi and includes targeted gene deletion and disruption by genetic engineering tools. The transfer of DNA constructs into fungal cells by genetic transformation with subsequent integration into the target locus via homologous recombination is an approach for gene inactivation [5]. Although there are several efficient methods for transformation of *T. atroviride* available [6-8], the generation of targeted gene deletion/disruption mutants is tedious and time-consuming, probably due to a low frequency of site-specific integration of the transforming DNA.

The recent adaption of the CRISPR/Cas9 system for fungi has significantly facilitated targeted gene-editing and genetic manipulation in these organisms. This is also true for *Trichoderma* spp., among which the industrial enzyme producer *T. reesei* was one of the first fungal species for which this technology has been employed [9, 10]. CRISPR/Cas9-mediated gene-editing is based on the introduction of DNA double-strand breaks (DSBs) at defined sites in the genome by the Cas9 endonuclease that is guided to the target locus by either a respective single guide RNA (sgRNA) or a duplex guide RNA (gRNA) formed by combining together the custom-designed CRISPR-derived RNA (crRNA) and the trans-activating CRISPR RNA (tracr RNA) [11]. Besides a 20 bp region in the sgRNA being homologous to the target site in the genome, *Streptococcus pyrogenes*-derived Cas9 needs a so-called protospacer adjacent motif (PAM) to be present 3’ of the target site [12, 13]. For application in gene-editing, both, Cas9 and sgRNA(s) have to be delivered into the fungal cell. This can be achieved by transforming and expressing plasmid-encoded Cas9 and sgRNA(s) in *vivo*, by combining *in vivo* expressed Cas9 with *in vitro* generated sgRNA(s)/gRNA(s), or by transforming an *in vitro* pre-assembled ribonucleoprotein (RNP) complex consisting of Cas9 and sgRNA(s)/gRNA(s) [10]. While long-term expression of Cas9 in the cell may cause unwanted, *e*.*g*. off-target, effects, the latter approach allows an only transient activity of the endonuclease.

Repair of the Cas9-generated DNA double-strand breaks may occur through different DNA repair mechanisms present in the target cell [14]. In most fungi, the error-prone non-homologous end-joining (NHEJ) pathway dominates over template-dependent homologous recombination (HR) and mainly results in the introduction of random mutations such as deletions or insertions at the target site. Tailor-made and precise editing is the result of HR, but requires the delivery of a repair template harboring sequences homologous to the flanks of the break [15, 16]. Several studies showed that activation of HR for repair of the Cas9-generated DNA double-strand breaks can be achieved even with short homology flanks on the repair template, while long homologous regions of about 1 kb are required with conventional gene-editing approaches [17-20].

Although the CRISPR/Cas9 system is highly efficient, markers are still needed for transformant selection. In *Trichoderma* spp., *hph* (mediating resistance to the antibiotic hygromycin B) and *pyr4* (auxotrophic marker encoding orotidine 5-phosphate decarboxylase) are most frequently used for this purpose [7]. Strains deficient in *pyr4* require the provision of uridine/uracil and are resistant to 5-fluoroorotic acid (5-FOA) which otherwise is metabolized to toxic fluorouracil by orotidine 5-phosphate decarboxylase. The *pyr4* gene hence is suitable for auxotrophic selection and 5-FOA counterselection. Nevertheless, markers for *T. atroviride* transformation are limited causing the need for marker recycling for the generation of mutants requiring sequential genetic manipulations. One strategy to address this challenge is to avoid integration of the selectable marker into the fungal genome, for example when using autonomously replicating AMA1-containing plasmids. They do not integrate into the genome and get lost upon repeated growth on non-selective media [21, 22]. A similar principle is represented by transiently selectable telomere (pTEL) vectors which contain a pair of telomeres and in the fungal cell behave as artificial chromosomes. In fungi, this kind of vector was first used in *Podospora anserina* for cloning purposes and has recently been combined with the CRISPR/Cas9 system for marker-free genome editing in *Botrytis cinerea* with a rapid loss of resistance during growth on non-selective medium [17, 23]. By using such vectors, the encoded selection markers are recyclable and can be used repeatedly for sequential rounds of transformation into the same recipient strain.

In this study, we show the successful and efficient genetic manipulation of *T. atroviride* strain P1 by transformation of fungal protoplasts with pre-assembled Cas9-gRNA RNPs. We established the *pyr4* gene as an additional selection marker for *T. atroviride* P1 by generating uridine/uracil auxotrophic *pyr4* disruption mutants by triggering a single Cas9-mediated DSB followed by NHEJ-based repair. The resulting mutants were used as recipients for transformation with a *Neurospora crassa pyr4* marker gene fragment in order to prove the functionality of the selection system. Furthermore, transformant selection via transiently stable telomeric vectors in *T. atroviride* was established. By combining pTEL-hyg transformation with two Cas9-gRNA RNPs aiming at excision of the target gene, marker-free *pks4*-deficient mutants impaired in conidial pigment synthesis as well as mutants devoid of the peroxin-encoding *pex5* gene were generated.

## 3 Materials and methods

### 3.1 Strains and culture conditions

*T. atroviride* strain P1 (ATCC 74058) [24] was used in this study and is referred to as the wild type (wt). Fungal strains (Supplementary Table S1) were cultivated and maintained on potato dextrose agar (PDA, Difco) or *Trichoderma* minimal medium (TMM) at 25 °C. The recipe for TMM was adapted from [25] without the addition of peptone and implementing the trace elements according to Cove’s recipe [26]. Fungal conidia were suspended in a solution with 0.9% NaCl and 0.01% Tween80. All bacterial plasmids were propagated in *Escherichia coli* Stellar^TM^ cells (Takara) using 100 µg/mL ampicillin or 50 µg/mL kanamycin for selection.

### 3.2 crRNA design and ribonucleoprotein (RNP) assembly

crRNA, tracrRNA, and the Cas9 protein, were purchased from Integrated DNA Technologies (IDT, Newark, USA). To design the specific crRNAs, IDT internal custom gRNA tool (https://eu.idtdna.com/site/order/designtool/index/CRISPR_CUSTOM) and ChopChop (https://chopchop.cbu.uib.no/) were used. To prevent unintended off-target DNA editing, the proposed gRNA candidates were used in a Blastn search against the *T. atroviride* P1 genome (https://mycocosm.jgi.doe.gov/Triatrov1/Triatrov1.home.html). Candidates showing an off-target sequence with more than 15 bp, including a PAM-site or minus (non-coding) strand target, were eliminated. Duplex formation of crRNA and tracRNA was carried out according to the IDT manufacturer’s specifications. RNP complexes were assembled containing 27 µg Cas9 enzyme (2.7 µL), 1.5 µL 10x Cas9 buffer, 7 µL gRNA (duplex, 20 µM) and 3.8 µL nuclease free water, resulting in an end volume of 15 µL and a molar ratio of 1:1.2 of RNA to Cas9 enzyme. The 10x Cas9 buffer (200 mM HEPES, 1.5 M KCl, 82 mM MgSO_4_x7H_2_0, 1mM EDTA and 5mM DTT at pH 7.5) was adapted from [27]. The RNP mixture was incubated at 37 °C for 15 min and prepared freshly before transformation.

### 3.3 Generation of transformation constructs, transformants selection and verification

As a reference genome, the *T. atroviride* strain P1 genome from the MycoCosm portal (https://mycocosm.jgi.doe.gov/Triatrov1/Triatrov1.home.html) was used. Genome architecture into distinct chromosomes was determined based on the information provided by the genome assembly ASM2064779v1 (https://www.ncbi.nlm.nih.gov/datasets/genome/GCF_020647795.1/ [28]. All primers and crRNA sequences used in this study are listed in Supplementary Table S2. To disrupt *T. atroviride pyr4* orthologue (Triatrov1 43450), the single RNP guide crPyr4_5’ downstream, located close to the start codon, was used. TMM containing 1.5 mg/mL 5-FOA, 10mM uridine and 5mM uracil was employed for the selection of the transformed cells. *pyr4* mutant candidates were verified through conventional PCR (primer pair pyr4_1.1/pyr4_2.1) and Sanger sequencing using the primer pyr4_seq1 (Microsynth AG, Balgach, Switzerland).

For deletion of the *T. atroviride pks4* gene (Triatrov1 488570) with the help of a recyclable marker, CRISPR assisted transformation was performed by utilizing dual 5’ and 3’ RNPs (crPks4_5’and crPks4_3’) to introduce DSBs at the beginning and at the end of the coding region. RNP-complexes (27 µg Cas9, each) were co-transformed with 10 µg pTEL-hyg [17] into *T. atroviride* protoplasts. For selection, 200 µg/mL hygromycin B was used in the PDA transformation plates. Transformants with white conidia were isolated and examined for gene editing events at the *pks4* gene locus by PCR genotyping using primer pairs pks4_2.3/pks4_1.4, pks4_1.4/ pks4_2.4 and pks4_1.3/pks4_2.3. For analysing the presence or absence of pTEL-hyg in the primary transformants and their progeny, primer pair pTEL_cpc1-hygR_F1 /pTEL_cpc1-hygR_ R1 was used.

Deletion of *pex5* gene (Triatrov1 428665) was performed in both *T. atroviride* wt and *pyr4* mutant background. In both cases, the two Cas9 RNPs were guided by the crRNAs crPex5_5’ and crPex5_3’. For the wt strain, 10 µg of pTEL-hyg was co-transformed for selection, while in the *pyr4* mutant (mutant 10.2), the *pex5* gene was replaced by a classical HR template carrying a functional copy of the *N. crassa pyr4* gene (NcPyr4) between homologous *pex5* flanks of about 1-kb. The 1779-bp NcPyr4 fragment was amplified from *N. crassa* genomic DNA [29] using the primer pair Nc_pyr4_fwd Nc_pyr4_rev. The homologous flanks were amplified from *T. atroviride* wt gDNA: the 812-bp 5’ fragment with the primer pair 5’flank_pex5_fwd/5’flank_pex5_rev and the 1000-bp 3’ flank with the primer pair 3’flank_pex5_fwd/3’flank_pex5_rev. The three fragments were assembled using the NEBuilder® HiFi DNA Assembly Master Mix (New England Biolabs Inc., Ipswich, USA) and subsequently amplified with the primers 5’flank_pex5_fwd/3’flank_pex5_rev and subcloned into pJET 1.2/blunt vector using the CloneJET PCR Cloning Kit (Thermo Scientific Inc., Waltham, USA). The plasmid was designated as pKOPex5 and verified by Sanger sequencing with primers pJET 1.2 forward seq and pJET 1.2 reverse seq. 4 µg of the HR fragment generated by PCR with primers 5’flank_pex5_fwd/3’flank_pex5_rev were used for transformation.

Transformants were utilized for DNA extraction and initially analysed through conventional PCR to verify genome editing, using primer pex5_1.1 and pex5_2.1 for gene excision in the wt, followed by Sanger sequencing for additional confirmation. For *pyr4* mutant background primer pex5_1.1 together with Nc_pyr4_2.3 and Nc_pyr4_1.3 together with pex5_2.1 were used for confirmation of gene replacement.

### 3.4 CRISPR/Cas9-assisted transformation of *T. atroviride*

As starting material, 1 × 10^6^ conidia/mL (final concentration) were inoculated into two 200 mL Erlenmeyer flasks containing 50 mL TMM and cultivated for 16-20 h at 25°C with shaking at 250 rpm. Mycelia were harvested aseptically by filtration, washed with sterile water, resuspended in 15 mL fungal lysis buffer (4% [w/ vol] VinoTastePro lytic enzyme Novozymes mix, 0.6 M KCl, 25mM potassium phosphate buffer, pH 5.8) and incubated at 30°C with shaking at 75 rpm until protoplasts emerged (approximately 2 to 4 hours). The digestion mixture was filtered through sterile filter papers (4 – 12 µm pore size) to get rid of cell wall debris and protoplasts were harvested by centrifuging for 15 min at 1,200 g and 4°C. Protoplasts were then washed twice with 0.6 M KCl and centrifuged. The resulting pellet was finally resuspended in 175 µL STC buffer (1 M sorbitol, 50 mM CaCl_2_, 10 mM Tris-HCl, pH 7.5) with a final concentration of about 5 x 10^6^ protoplasts/mL.

The transformation procedure for the RNP complex was adapted from Kwon *et al*. [30] and Pohl *et al*. [27]. For each Cas9-mediated transformation, 150 μL protoplasts, 5 μL HR donor DNA (5 μg) if indicated, 15 μL RNP complex, 20 μL 2× STC, 20 μL 10 × Cas9 buffer and 50 μL 60 % PEG 4000 buffer (50 mM CaCl2 and 10 mM Tris-HCl, pH 7.5) were mixed in a 50 mL tube. The mixture was incubated on ice for 25 min, followed by a 2-minute equilibration to room temperature. Afterwards, 1 mL of 60 % PEG 4000 was added, followed by an incubation step of 20 min at room temperature. Subsequently, the mixture was diluted with 5 mL STC buffer and filled up to 25 mL with the appropriate top agar medium (0.9 % agar supplemented with 1 M sorbitol) with the respective selection condition. The solution was equally distributed onto five plates with pre-poured bottom media supplemented with the respective selection condition. Plates were incubated at 25° C for 4-5 days, emerging transformants transferred to fresh plates and subsequently subjected to three rounds of single spore isolation in order to obtain homokaryotic strains.

### 3.5 Biotin auxotrophy testing

10^4^ conidia were point inoculated on TMM, prepared either without or supplemented with biotin (1µg/ mL). Colony diameters were measured in triplicate after five days. Results were validated with an ANOVA statistical analysis with post-hoc Tukey HSD to assess the significance of differences.

## 4 Results

### 4.1 Establishing CRISPR/Cas9-assisted transformation in *T. atroviride* while creating a new selectable marker

CRISPR/Cas9-based gene inactivation relying on repair of the generated DSBs through non-homologous end joining (NHEJ) has successfully been applied in diverse fungi, including *T. reesei* [9, 31], but was so far not tested in *T. atroviride*. To assess whether a single Cas9-mediated DSB induced by pre-assembled Cas9-sgRNA RNP could be suitable for gene disruption in *T. atroviride*, we selected the putative homolog of *pyr4* as target. The *pyr4* gene is a commonly used selection marker in various fungal species [32, 33]; however, no respective *pyr4* mutant of *T. atroviride* P1 has been available so far. BLASTp analysis resulted in the identification of Triatrov1 34450 as the *T. atroviride* P1 orotidine 5’-phosphate decarboxylase, encoded by the *pyr4* gene orthologue, sharing 89% and 60% amino acid identity with Pyr4 proteins from *T. reesei* and *N. crassa*, respectively. For generating *pyr4-*deficient mutants in *T. atroviride* P1, a single gRNA was designed to direct Cas9 cleavage 210-bp downstream of the start codon (Figure 1).

**Figure 1.**
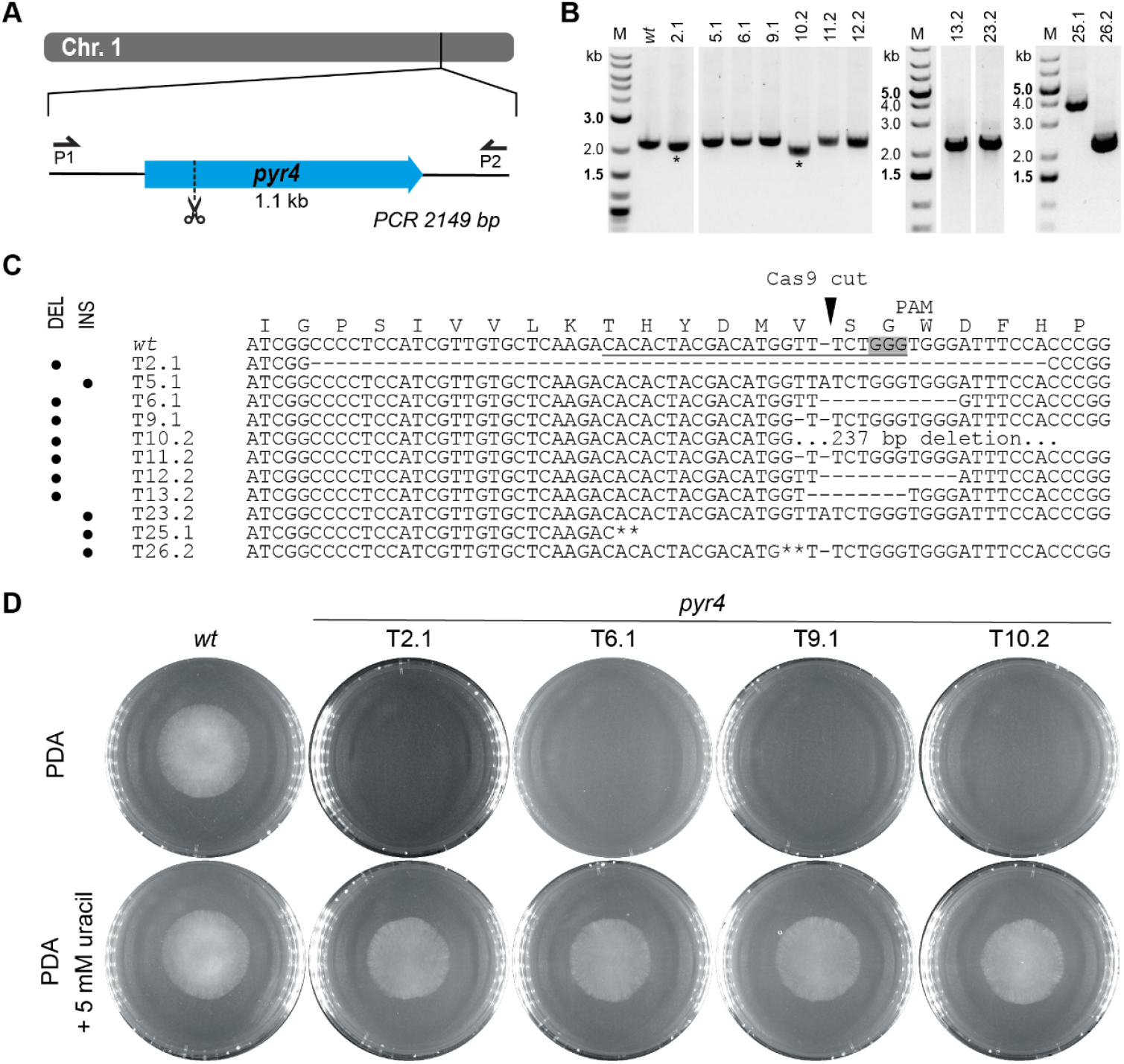
Inactivation of the *pyr4* gene in *T. atroviride* P1. (A) Schematic diagram of the *pyr4* coding sequence on chromosome 1 with the Cas9 target site indicated scissors. Primers P1 (pyr4_1.1) and P2 (pyr4_2.1) were used for locus amplification. (B) Genotyping of eleven selected 5-FOA resistant and uracil/uridine auxotrophic transformants and *T. atroviride* wt. Lanes marked with an asterisk (*) indicate transformants with deletions in the *pyr4* locus. (C) Amplicon sequencing results covering a part of the *pyr4* locus of the wt and eleven selected transformants. The corresponding amino acid sequence is given above the genomic sequence. The region targeted by the gRNA is underlined, the PAM sequence is framed in grey and the Cas9 cutting site is marked with a triangle. Asterisks (**) indicate the integration of sequences from different chromosomal regions. DEL indicates mutants with deletions in the py4 locus; INS indicates mutants with insertions. (D) Comparative assessment of growth of the wt and four *pyr4* mutants on either PDA or PDA supplemented with 5 mM uracil.

Inactivation of *pyr4* is expected to result in uridine/uracil auxotrophy and resistance to 5-FOA. 163 transformants displaying 5-FOA resistance were obtained, indicating a loss-of-function mutation in the *pyr4* gene. Of these, 26 transformants were randomly chosen and tested for growth on a minimal medium deficient in uracil or uridine. Except for one transformant that exhibited weak growth, none of the mutants were able to grow unless the medium was supplemented with 5 mM uracil or 10 mM uridine.

11 transformants were randomly selected and subjected to sequencing, which revealed alterations in the mutants *pyr4* coding sequence as result of error-prone repair of the DSB by NHEJ. In seven transformants (2.1, 6.1, 9.1., 10.2, 11.2, 12.2, 13.2) deletions from 1-bp up to 237-bp had occurred at the Cas9 target site, in two mutants (5.1 and 23.2) an adenine-containing base was inserted, while mutants 25. 1 and 26.1 showed insertions of sequences with homology to other chromosomal regions or even a different chromosome, respectively. Transformant 26.1 exhibited at the Cas9 cutting site an 86-bp insertion from chromosome 2, while an insertion of about 1.7-kb, containing sequences from chromosome 1, had occurred in transformant 25.1 (Figure 1 C).

### 4.2 Using transient telomeric vectors for marker recycling in combination with CRISPR/Cas9

For generation of *T. atroviride* mutants without any integrated resistance markers, we applied two Cas9-sgRNA RNPs targeting different regions in the *pks4* gene locus, thereby aiming at excision of the coding sequence between the generated DSBs by NHEJ (Figure 2 A). As previously shown in *T. reesei*, Pks4 is responsible for the biosynthesis of the conidial pigment and Δ*pks4* mutants can easily be identified morphologically by their unpigmented conidia [34], enabling a straightforward evaluation of *pks4* gene deletion in the resulting transformants. The *T. atroviride* P1 Pks4 orthologue (ID488570) was identified by BLASTp analysis with *T. reesei* Pks4 (XP_006969537) and *A. fumigatus* PksP (XP_756095) as queries with which it showed a sequence identity of 72.4% and 51.9%, respectively. It is important to note that *T. atroviride pks4* was found to be located in the subtelomeric region of chromosome 5 with the distance to the end of the chromosome being about 13 times the length of the *pks4* coding region.

**Figure 2.**
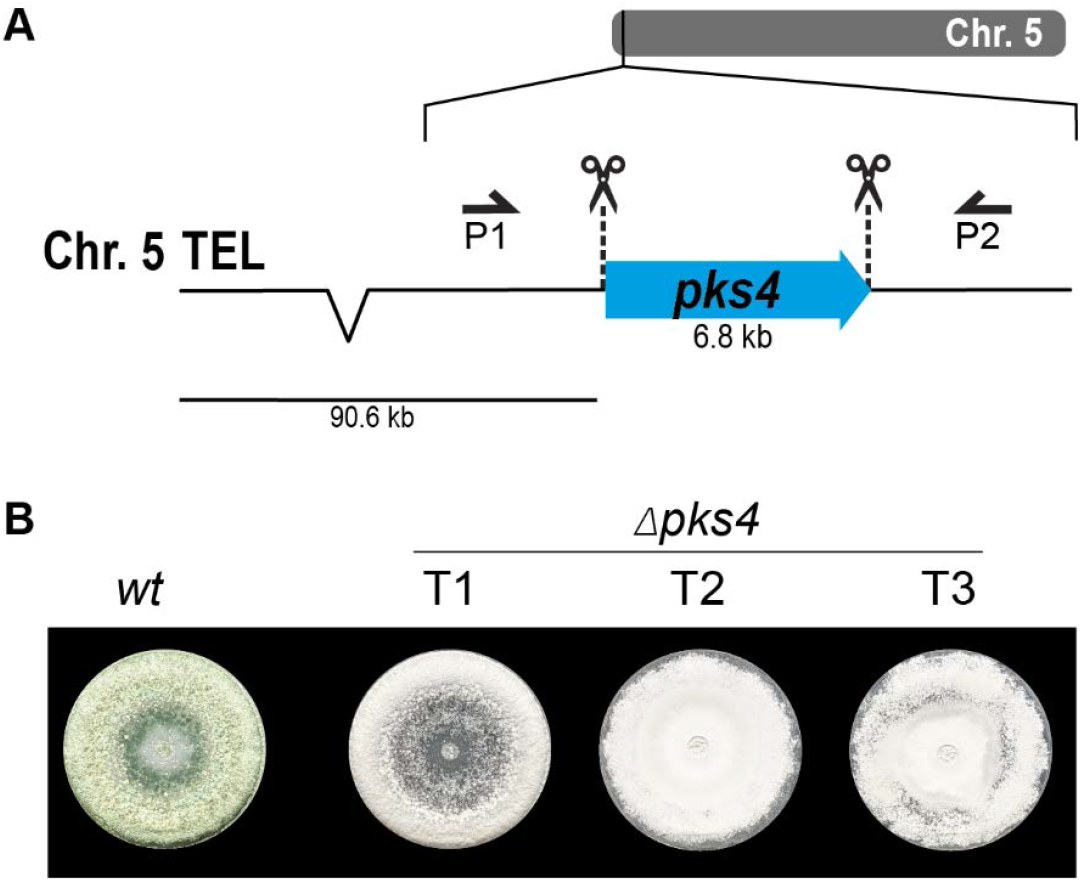
Deletion of *pks4* through a bipartite marker-free editing approach. (A) Schematic diagram of the *pks4* coding sequence in the subtelomeric region of chromosome 5 with the Cas9 target sites indicated by scissors. Primers P1 (pks4_1.4) and P2 (pks4_2.3) were used for locus amplification. (B) Absence of conidial pigmentation being indicative of *pks4* gene inactivation in transformants 1, 2 and 3 (T1, T2, T3) upon growth on PDA.

For selection, fungal protoplasts were co-transformed with pTEL-hyg [17], a plasmid harbouring telomeric sequences and the *hph* resistance cassette. Co-transformation yielded a total of 182 colonies. From these, 38 transformants were randomly selected and further cultured. Of these, only nine displayed the wild-type green spore pigmentation, while the remaining 29 (76%) produced unpigmented white spores (Figure 2 B). One green-pigmented and seven non-pigmented transformants were selected for PCR-based genotyping. We could detect an alteration in the *pks4* locus for all seven white strains compared to the wild type and the green conidia producing transformants, but we failed to verify the deletion of the complete gene. Despite the white spore phenotype, a smaller fragment of the coding region could be amplified in some candidates, indicating that the *pks4* sequence was not completely lost.

pTEL vectors and their associated resistance markers are rapidly lost in *B. cinerea* and *Magnaporthe oryzae* upon fungal cultivation on non-selective medium [17]. Obtained *T. atroviride pks4* mutant strains hence were transferred to PDA without hygromycin B and after each passaging, the new generation of conidia was tested for loss of antibiotic resistance. Only two out of seven tested mutants became hygromycin-sensitive in the first and tenth generation, respectively. Vector loss was further confirmed on the genetic level; no pTEL-hyg sequences could be amplified from the mutants via PCR. Although the pTEL vector did not get lost as efficiently and quickly as previously reported for *B. cinerea* and *M. oryzae*, the presence of numerous white transformants confirms the generation of the intended changes in the *pks4* locus by the transformed Cas9-gRNA RNPs thereby resulting in *pks4* loss-of-function mutants.

### 4.3 Comparison of the two newly developed marker strategies in *T. atroviride*

To prove the applicability of the developed approaches in *T. atroviride*, we selected the *pex5* gene as new target. Marker-free deletion mutants were generated through co-transformation of Cas9-sgRNA RNPs either with *pyr4* as new selectable marker or with the pTEL-hyg vector. *pex5* encodes the peroxisomal targeting signal type-1 (PTS1) receptor, a cytosolic cycling adaptor protein required for protein import into the matrix of peroxisomes [35-37]. The *T. atroviride* Pex5 orthologue (Triatrov1 428665) was identified by BlastP analysis using *Asperillus nidulans* PexE (XP_659120) and *N. crassa* Pex5 (XP_965347) as queries, with which it showed an amino acid sequence identity of 62.6 % and 54.3 %, respectively [36, 38].

For the first approach, a classical gene replacement strategy was used by applying the *pyr4* marker system for selection. To this end, the above-described *T. atroviride pyr4-*deficient recipient was transformed with a functional fragment of the *N. crassa pyr4* gene, including its endogenous promotor and terminator [39], flanked by about 1 kb homology region to *pex5* upstream and downstream sequences for homologous recombination (Figure 3 A). For the second approach, T. *atroviride* was co-transformed with two Cas9-sgRNA RNPs and the pTEL-hyg vector for selection aiming at excision of the *pex5* ORF by generating two DSBs via the CRISPR/Cas9 system followed by NHEJ-based repair (Figure 4 A). By using these two approaches, the efficiency of gene knockout by NHEJ or HDR targeting the same gene with identical RNPs could be compared.

**Figure 3.**
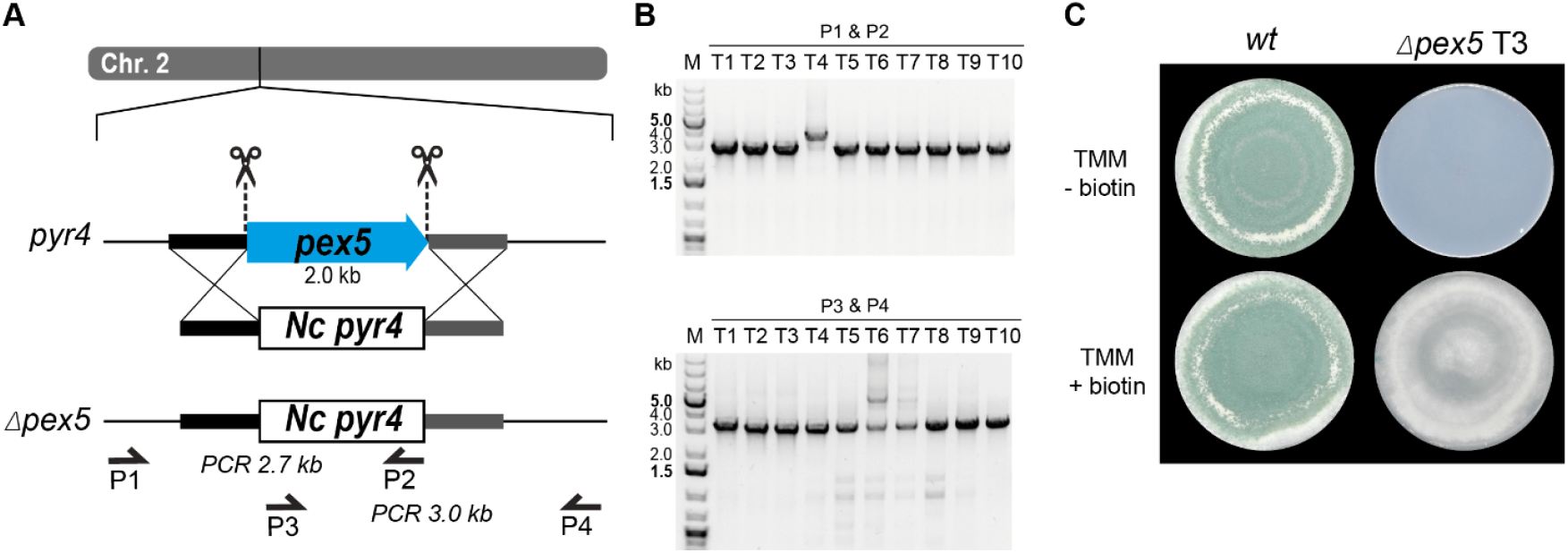
Deletion of *pex5* through a CRISPR/Cas9 assisted gene replacement approach in Δ*pyr4* background. (A) Schematic diagram of the *pex5* coding sequence on chromosome 2 with the Cas9 target sites indicated by scissors. Primers P1 (pex5_1.1), P2 (Nc_pyr4_2.3) and P3 (Nc_pyr4_1.3), P4 (pex5_2.1) were used for locus amplification. *Pex5* is replaced by the *N. crassa pyr4* gene. (B) Genotyping of ten transformants using primer pair P1, P2 (top) or P3, P4 (bottom). (C) Comparative assessment of growth of the wt and transformant 3 (T3) on either TMM or TMM supplemented with 1µg/mL biotin.

**Figure 4.**
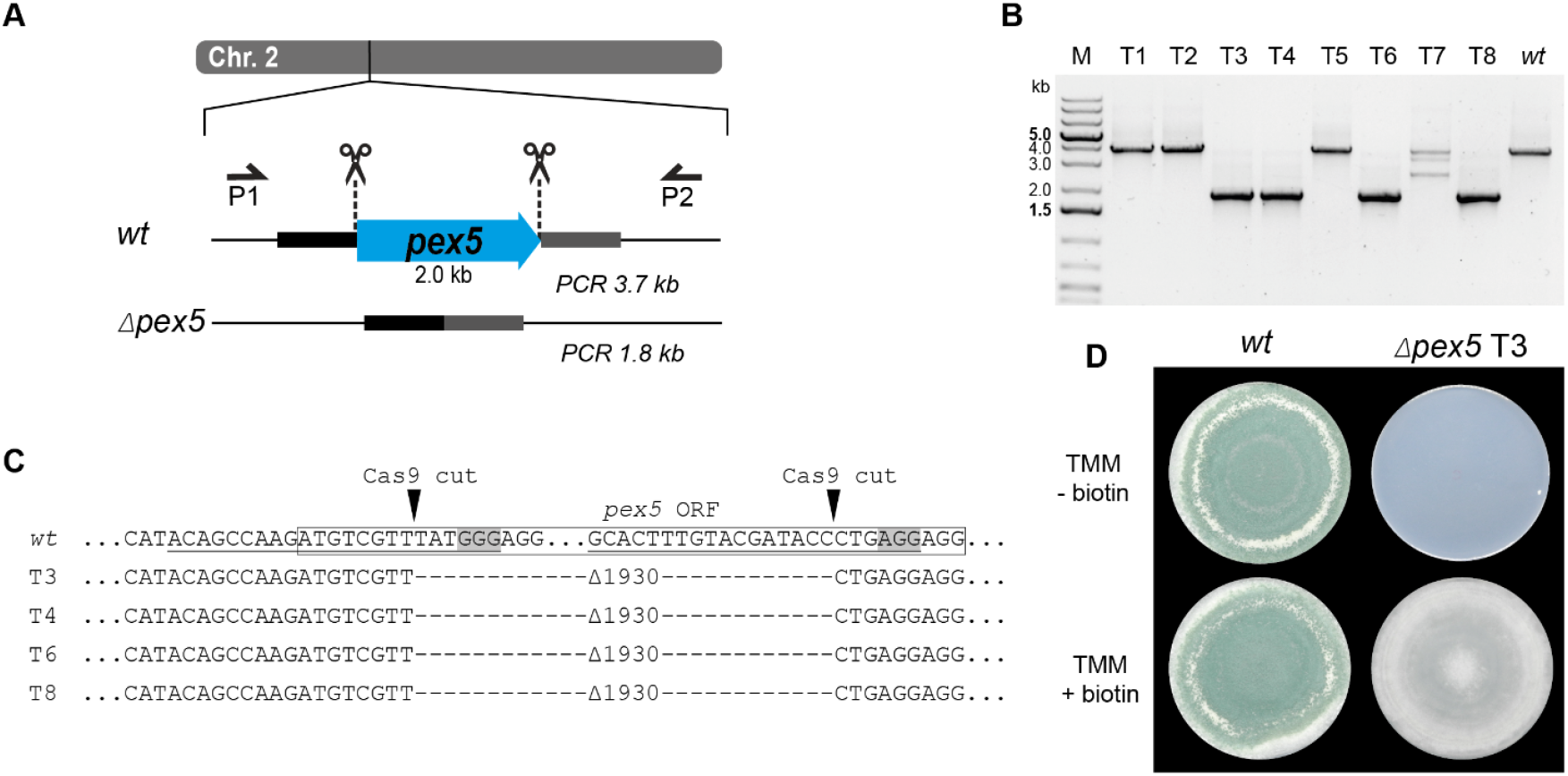
Deletion of *pex5* through CRISPR/Cas9-mediated excision. (A) Schematic diagram of the *pex5* coding sequence on chromosome 2 with the Cas9 target sites indicated by scissors. Primers P1 (pex5_1.1) and P2 (pex5_2.1) were used for locus amplification. Excision of the *pex5* coding sequence is followed by non-homologous end-joining (NHEJ), resulting in Δ*pex5* mutants. (B) Genotyping of eight transformants and *T. atroviride* wild type (wt). (C) Amplicon sequencing results covering part of the *pex5* locus of the wt and transformants 3, 4, 6 and 8 (T3, T4, T6, T8). The region targeted by the gRNA is underlined, the PAM sequence is framed in grey and the Cas9 cutting site is marked with a triangle. The *pex5* open reading frame (ORF) is enclosed in a rectangle. A DNA fragment of 1930 bp was excised. (D) Comparative assessment of growth of the wt and transformant 3 on either TMM or TMM supplemented with 1µg/mL biotin.

Transformation using the *pyr4* selection marker system resulted in a total of 237 uridine/uracil prototrophic transformants, confirming successful complementation by the *N. crassa pyr4* gene. Ten randomly selected transformants were genotyped by PCR to verify successful replacement of the *pex5* target locus by *N. crassa pyr4* (Figure 3 B).

Use of the pTEL-hyg vector for selection resulted in 64 transformants. PCR-based genotyping and *pex5* locus sequencing of eight randomly selected transformants confirmed deletions at the target locus in four of them (Figure 4 B). After cultivation on non-selective media, the tested transformants lost hygromycin B resistance within two generations.Loss of *pex5* resulted in impaired conidiation and a biotin auxotrophic phenotype, which is consistent with mutants of other fungi with non-functional PTS1 import (Figure 3 C; Figure 4 D) [38, 40]. Interestingly, biotin supplementation could not fully restore radial colony growth in the generated *T. atroviride Δpex5* mutants to the level of the wild type, regardless of whether the mutants derived from the wild type or the *pyr4-*deficient recipient as parental strain (data not shown).

## 5 Discussion

In this work, we developed two strategies to overcome the problem of selectable marker availability in *T. atroviride*. First, we generated an uracil/uridine auxotrophic strain that can be used as recipient for genome editing experiments, employing the *pyr4* gene from *N. crassa* for complementation and selection. Then we proved the applicability in *T. atroviride* of a telomeric vector that can be used in a co-transformation approach and allows recycling of the hygromycin marker cassette. Both strategies were achieved while implementing protoplast-based transformation with the use of the CRISPR/Cas9 system delivered as RNP. RNP-mediated CRISPR/Cas9 genome editing is more effective and avoids the problems associated with using Cas9-encoding plasmids, as the use of RNPs reduces total experimental time. Moreover, this approach limits Cas9 off-target effects, as RNPs are present only transiently in the cell while the plasmid-harbored *cas9* gene normally has to integrate into the host genome [41, 42]. Furthermore, the potential toxicity of Cas9 in fungal cells restricts applications with stable Cas9 expression[43, 44].

The delivery of RNP complexes must be specifically adapted and optimized for each organism, and their suitability thoroughly tested to ensure effective application. Here we demonstrated the successful application of CRISPR/Cas9 RNPs for gene deletion or disruption in *T. atroviride* with three different targets and approaches.

The *T. atroviride pyr4* gene (Triatrov1 34450) encodes the orotidine 5’-monophosphate decarboxylase, which is involved in the pyrimidine biosynthetic pathway [33, 45]. Deletion of *pyr4* is known to generate uracil/uridine auxotrophy but, at the same time, to confer resistance to 5-FOA [46]. We successfully managed to generate several *pyr4-*deficient strains employing a single RNP that targeted the 5’ region of *pyr4* coding sequence. These auxotrophic mutants were the result of a single DSB and NHEJ repair mechanism which, according to our sequencing results, generated various types of loss of function mutations, including different size deletion, frame shift and even insertions of several bases. Similar results were obtained with a biolistic plasmid-based Cas9 and gRNA approach for genome editing in *Trichoderma harzianum*, although with fewer transformants [47]. Unexpected genome editing, where larger fragments were integrated during NHEJ, has already been observed in *T. reseei* targeting the orotate phosphoribosyl transferase encoded by *ura5*, which also led to uridine/uracil auxotrophy [31]. The results from this experiment suggests that our protocol for RNP delivery into *T. atroviride* is effective and efficient, and can be considered a valuable upgrade of the classical protoplast-based transformation approach. The efficiency of transformation with single RNP was considered more than satisfying even without providing a HR oligo for repair of the DSB and also without supplementation of Triton X-100 [20].

An even more successful gene deletion was achieved by co-transforming the RNPs with a transiently stable telomere vector containing the *hph* gene, which was rapidly lost when cells were grown on non-selective media, following a strategy previously used in *B. cinerea* [17]. For improved gene editing, the protocol was modified to generate deletions using two RNPs targeting the same gene, without the addition of a repair template. This modification resulted in the excision of the sequence between the cleavage sites through the more efficient NHEJ repair mechanism. The efficiency of this strategy was first tested using *pks4* as target gene, which encodes a polyketide synthase responsible for conidial pigmentation [34, 48, 49]. The use of such target allowed us to quickly identify positive transformants by the lack of green pigmentation of their spores. Several white colonies grew on the transformation plates, many of which were selected for genotyping verification. Determination of the kind of mutations that occurred between the two RNP target sites, however, proved challenging as removal of such a large gene (6.8 kb) might lead to a variety of recombination events. In addition to this newly established approach, recent studies have shown that the AMA plasmid can be used for marker recycling in *T. reesei* with CRISPR/Cas9 gene editing, making it a potential alternative to the pTEL vector in *T. atroviride* as well [50].

As proof of concept, to validate the efficacy of our developed approaches, we selected *pex5* (Triatrov1 428665) as target for deletion, which encodes the peroxisomal targeting signal type-1 (PTS1) receptor, a cytosolic cycling adaptor protein required for protein import into the matrix of peroxisomes [36, 38]. Both strategies, replacement of *pex5* with a functional copy of *pyr4* from *N. crassa* by HR or excision of *pex5* with NHEJ repair in co-transformation with pTEL, were found extremely efficient, generating a large number of mutant strains. The co-transformation with the transiently stable vector, in particular, led to four marker-free strains with the desired editing out of eight transformants tested after a single transformation attempt.

Our experiments with *pex5* as a target showed precise excision by CRISPR/Cas9 RNP and error-free sealing of the gap by NHEJ, similarly to what has been reported by Leisen *et al*. in *B. cinerea* [17, 51]. We also have to emphasize that NHEJ is not error-prone by default, as often mentioned in respect to DSB repair in gene editing. This would be fatal for fungi predominately using NHEJ over HDR for DNA repair [52] and NHEJ is inherently accurate in the repair of Cas9-induced DSBs, leading to a high frequency of accurate NHEJ events [53]. In addition, it was observed in *Aspergillus niger* that the CRISPR/Cas9-mediated double-strand break can be simultaneously repaired with a template DNA by the NHEJ and HDR pathways [15]. Thus, it is not fully understood how a given fungal species responds to a Cas9-induced DSB for gene editing and which repair mechanisms occur in which order or even simultaneously. As the selection process is independent of the genome editing process, transformants containing the pTel vector but without gene deletion were expected. However, a sufficient number of transformants exhibited the desired excision and, to our surprise, were immediately homokaryotic, which is unusual for *Trichoderma* species.

In conclusion, our streamlined approach—using CRISPR/Cas9 RNP delivery in combination with transiently stable telomere vectors—enables efficient, marker-free gene deletion and marker recycling in *T. atroviride* without the need for laborious knock-out cassette construction. By leveraging NHEJ over the less efficient HDR pathway, we have developed a versatile and time-saving reverse genetics tool. This system not only accelerates functional gene analysis but also strengthens the genetic toolkit available for *Trichoderma* species, which hold growing importance in both basic research and biotechnological applications.

## 7 Author Contributions

MG: conceptualization, data curation, investigation, supervision, visualization, writing - original draft, writing - review and editing. CE: investigation. LD: investigation. US: investigation. SP: visualization, writing - original draft, writing - review and editing. CB: writing - original draft, writing - review and editing. SZ: funding acquisition, resources, writing - original draft, writing - review and editing.

## 9 Acknowledgments

We thank Prof. Matthias Hahn (RPTU, Kaiserslautern) for kindly providing the pTEL-hyg plasmid used in this work and his valuable advice regarding its application.

